# BGC-QDR: A Quantum-Assisted Framework for Biosynthetic Gene Cluster Discovery and Drug-Potential Ranking

**DOI:** 10.64898/2026.06.21.733574

**Authors:** Abhishek Mishra, Ansh Rai

**Affiliations:** Bhagwan Parshuram Institute of Technology, Delhi, India

**Keywords:** Biosynthetic gene clusters, environmental DNA, variational quantum classifier, drug discovery, metagenomics, PennyLane, bioinformatics

## Abstract

Biosynthetic gene clusters (BGCs) encode enzymatic pathways for natural products with pharmaceutical potential, yet prioritizing candidates from fragmented environmental DNA (eDNA) assemblies remains computationally challenging. We present BGC-QDR (Biosynthetic Gene Cluster Quantum Discovery and Ranking), an open-source pipeline that integrates input quality control, Prodigal ORF prediction, Pfam HMM domain annotation, rule-based BGC classification, MiBIG 4.0 novelty assessment, and variational quantum classifier (VQC) ranking via PennyLane. BGC-QDR is designed as a **quantum-assisted ranking framework** for biologically informed BGC prioritization, not as a claim of quantum computational advantage over classical machine learning.

We evaluate the pipeline on MiBIG 4.0 (2,636 annotated BGCs) using a 20-dimensional biosynthetic feature vector and stratified 10-fold cross-validation. The integrated VQC (6 qubits × 3 layers, 54 parameters) achieves accuracy of 0.789 ± 0.076 and ROC-AUC of 0.835 ± 0.057. Random Forest achieves the highest ROC-AUC (0.898 ± 0.032), followed by Logistic Regression (0.874 ± 0.020) and MLP (0.872 ± 0.024). Wilcoxon signed-rank tests on per-fold AUC scores show that VQC ROC-AUC is significantly lower than Random Forest (*p* = 0.0098) and Logistic Regression (*p* = 0.037) at α = 0.05, with no significant difference versus MLP (*p* = 0.064). Architecture ablation identifies 4 qubits × 3 layers as the best VQC configuration on hold-out validation (AUC = 0.737). Feature importance analysis highlights peptidyl carrier protein domains, cluster length, and module count as dominant predictors. BGC-QDR provides a reproducible, end-to-end workflow for eDNA-derived BGC discovery with integrated novelty scoring and quantum-assisted candidate ranking. The complete BGC-QDR source code, benchmark datasets, and reproduction instructions are publicly available at: Abhishekmishra2808/BGC-PIPELINE

## I. Introduction

Natural products derived from microbial biosynthetic pathways remain a major source of antibiotics, immunosuppressants, and anticancer agents. Biosynthetic gene clusters (BGCs), genomic loci encoding polyketide synthases (PKS), non-ribosomal peptide synthetases (NRPS), terpene cyclases, and associated tailoring enzymes, are the genetic basis of this chemical diversity. Culture-independent metagenomic sequencing of environmental DNA (eDNA) has expanded access to microbial biosynthetic diversity, but eDNA assemblies are fragmented, incomplete, and difficult to prioritize for experimental validation.

Existing BGC detection tools such as antiSMASH, DeepBGC, and PRISM excel at identifying biosynthetic loci in complete or near-complete genomes. However, they generally stop at detection and classification; they do not provide an integrated, novelty-aware ranking of candidates by predicted drug-discovery potential. Furthermore, ranking BGCs requires labeled reference data. MiBIG 4.0 provides 2,636 annotated BGCs with activity labels, enabling supervised learning, but the practical value of quantum machine learning (QML) for this tabular biosynthetic feature problem remains an open empirical question.

We introduce BGC-QDR, a unified computational pipeline for BGC discovery and drug-potential ranking from metagenomic FASTA input. The pipeline comprises seven stages: quality control, ORF calling, Pfam domain annotation, BGC classification, MiBIG novelty assessment, VQC ranking, and output of ranked candidates (Figure 1). **We explicitly frame BGC-QDR as a quantum-assisted ranking framework**: the VQC provides a biologically interpretable, hybrid quantum-classical scoring component integrated into a practical bioinformatics workflow. We do **not** claim quantum advantage over classical machine learning; our experiments show that ensemble classical models achieve higher ROC-AUC on MiBIG 4.0.

**Figure 1:**
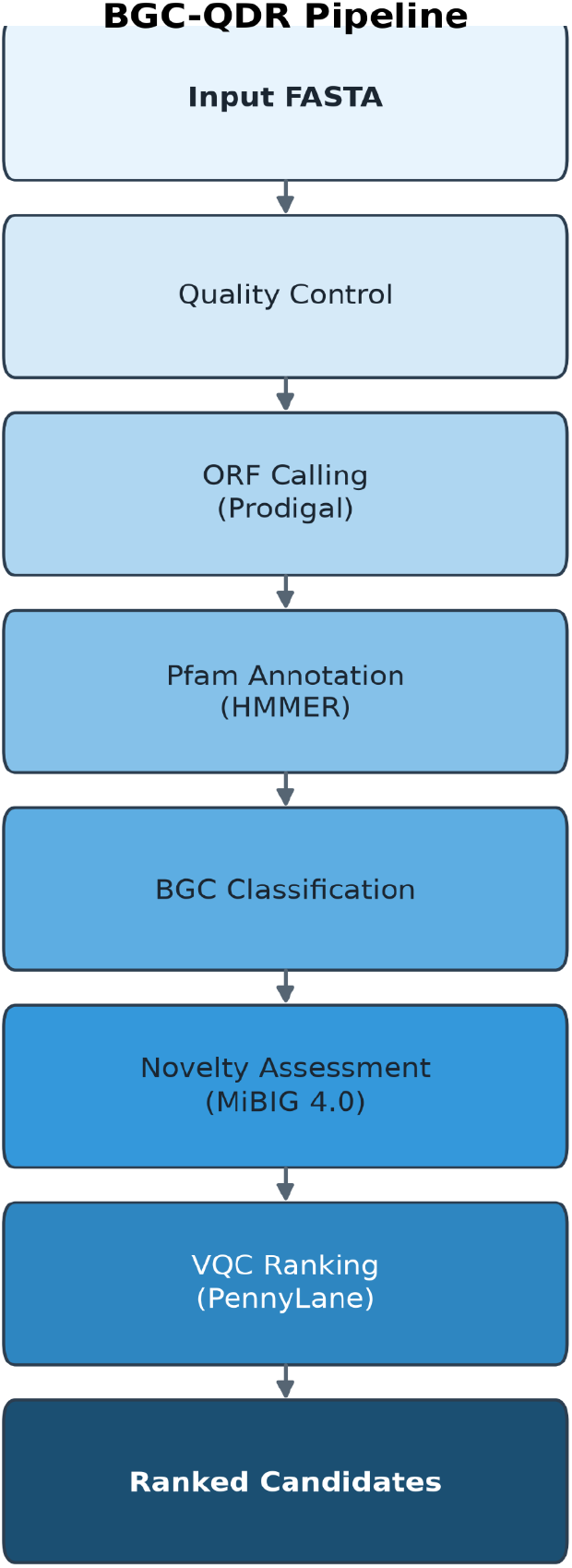
BGC-QDR end-to-end pipeline.

## Contributions

1. An open-source end-to-end pipeline for BGC detection, novelty scoring, and quantum-assisted ranking from eDNA or genomic FASTA without requiring complete genomes.
2. A 20-dimensional biosynthetic feature representation combining domain composition, cluster architecture, and biological diversity metrics, validated via Random Forest feature importance.
3. Rigorous benchmarking on MiBIG 4.0 with stratified 10-fold cross-validation, Wilcoxon signed-rank tests, and VQC architecture ablation (4 to 8 qubits, 2 to 3 layers).
4. A reproducible evaluation protocol with benchmark figures on MiBIG 4.0, a 15-locus antiSMASH comparison, and metagenomic tool benchmarking, showing that classical ensembles remain stronger rankers on ROC-AUC.

## II. Related Work

### BGC detection

Rule-based and HMM-driven tools (antiSMASH, PRISM) dominate reference-guided BGC annotation. Deep learning approaches (DeepBGC, SanntiS) improve sensitivity on novel architectures but typically require longer contigs. BGC-QDR adopts established domain-driven detection (Prodigal + Pfam + rule-based classification) for interpretability and compatibility with fragmented eDNA.

### Metagenomic BGC analysis

BiG-MAP and METABOLIC address abundance and screening at scale. BiG-SCAPE clusters BGCs into gene cluster families (GCFs) by domain similarity. BGC-QDR focuses on per-contig detection and ranking rather than large-scale GCF network analysis.

### Quantum machine learning

Variational quantum classifiers (VQCs) embed classical features into quantum circuits via angle encoding and trainable entangling layers. Empirical benchmarks consistently show that VQCs do not universally outperform classical models on tabular data; domain-specific evaluation is required. This work provides such an evaluation for BGC drug-potential ranking.

### ML-based natural product ranking

Domain architecture features predict bioactivity class in BGC contexts. BGC-QDR extends this with an integrated pipeline and comparative QML evaluation.

## III. Methods

### A. Pipeline Overview

Figure 1 illustrates the BGC-QDR workflow. Input metagenomic or genomic sequences (FASTA) pass through seven stages:

1. **Quality control:** filters short, low-complexity, and synthetic contigs.
2. **ORF calling:** Prodigal gene prediction on retained contigs.
3. **Pfam annotation:** HMMER domain scanning against Pfam-A (E-value ≤ 1×10^−5^).
4. **BGC classification:** rule-based typing from domain signatures (Type I PKS, NRPS, terpene, hybrid, and related classes).
5. **Novelty assessment:** domain-set Jaccard similarity against MiBIG 4.0.
6. **VQC ranking:** PennyLane variational quantum classifier trained on MiBIG features.
7. **Output:** ranked BGC candidates with quantum scores, novelty metrics, and pipeline log.

### B. Feature Representation

Each BGC is encoded as a **20-dimensional feature vector**:

- **15 domain-count features:** PKS, NRPS, terpene, condensation, AMP-binding, glycosyltransferase, P450, and related Pfam families.
- **2 structural features:** cluster length (kb), biosynthetic module count.
- **3 biological features:** domain Shannon entropy, resistance-gene proxy count, tailoring-enzyme count.

Features are Z-score normalized before model input. For the VQC, PCA reduces the 20 features to *n* principal components (where *n* equals the qubit count), scaled to [0, π] for angle embedding.

### C. Labels and Class Imbalance

MiBIG 4.0 drug-potential labels are binary (active vs inactive): **2**,**162 active (82.0%)** and **474 inactive (18.0%)**. This imbalance is expected because MiBIG prioritizes biochemically characterized, bioactive clusters.

We address imbalance at three levels:

1. **Splitting:** All cross-validation folds are **stratified** so each fold preserves the ~82/18 class ratio.
2. **Metrics:** We report **ROC-AUC as the primary metric** because it is threshold-independent and less sensitive to prevalence than accuracy alone. We also report precision, recall, and F1 for completeness.
3. **Models:** Logistic Regression and Random Forest use **balanced class weights**. The VQC is trained with **class-weighted binary cross-entropy**. The MLP uses early stopping on validation loss. Together, these settings reduce bias toward the majority (active) class while still allowing recall-focused behaviour where appropriate.

Because inactive BGCs are underrepresented, high recall can coincide with lower precision; we interpret accuracy and recall alongside ROC-AUC rather than relying on a single score.

### D. Variational Quantum Classifier

The VQC is implemented in PennyLane with the following default configuration (used in 10-fold CV):

- **Qubits / layers:** 6 qubits × 3 StronglyEntanglingLayers (54 trainable parameters).
- **Encoding:** AngleEmbedding (RY rotations) on PCA-reduced features.
- **Ansatz:** StronglyEntanglingLayers with ring-topology CNOT entanglement.
- **Measurement:** ⟨Z⟩ on qubit 0, mapped to class probability via sigmoid.
- **Training:** Adam optimizer (lr = 0.02), 60 epochs, batch size 5, class-weighted binary cross-entropy (see Section III.C).

Architecture ablation evaluates five configurations on hold-out data: 4q×2L, 4q×3L, 6q×2L, 6q×3L, 8q×2L.

### E. Classical Baselines

We compare against:

- **Logistic Regression:** linear separability baseline (class-weighted).
- **Random Forest:** 200 to 300 trees, balanced class weights.
- **MLP:** 128-64-32 hidden layers, early stopping.

All models receive the same 20-dimensional feature vectors with per-fold StandardScaler fitting.

### F. Evaluation Protocol and Statistical Testing

- **Dataset:** MiBIG 4.0 (2,636 BGCs; class distribution in Section III.C).
- **Cross-validation:** Stratified 10-fold CV; metrics reported as mean ± standard deviation across folds.
- **Primary metric:** ROC-AUC (class-imbalance robust; Section III.C).
- **Secondary metrics:** Accuracy, F1, precision, recall.
- **Pairwise tests:** Wilcoxon signed-rank test on per-fold ROC-AUC scores (VQC vs each classical model), α = 0.05.

With ten paired fold scores, the Wilcoxon test can reach conventional significance at α = 0.05 (minimum two-sided *p* = 0.002). We report both *p*-values and effect sizes (Table I, Figure 2); significant results indicate detectable differences in fold-wise ROC-AUC, not clinical or biological superiority in absolute terms.

**TABLE I.** Classification performance on MiBIG 4.0 (10-fold CV, mean ± std)

| Model | Accuracy | ROC-AUC | F1 | Precision | Recall |
| --- | --- | --- | --- | --- | --- |
| Random Forest | 0.759 ± 0.031 | **0.898** ± **0.032** | 0.832 ± 0.025 | 0.967 ± 0.013 | 0.731 ± 0.037 |
| MLP | **0.857** ± **0.015** | 0.872 ± 0.024 | **0.918** ± **0.010** | 0.867 ± 0.011 | **0.976** ± **0.024** |
| Logistic Regression | 0.726 ± 0.024 | 0.874 ± 0.020 | 0.805 ± 0.021 | 0.968 ± 0.012 | 0.689 ± 0.032 |
| **VQC (PennyLane)** | 0.789 ± 0.076 | 0.835 ± 0.057 | 0.860 ± 0.067 | 0.910 ± 0.051 | 0.831 ± 0.134 |

**Figure 2:**
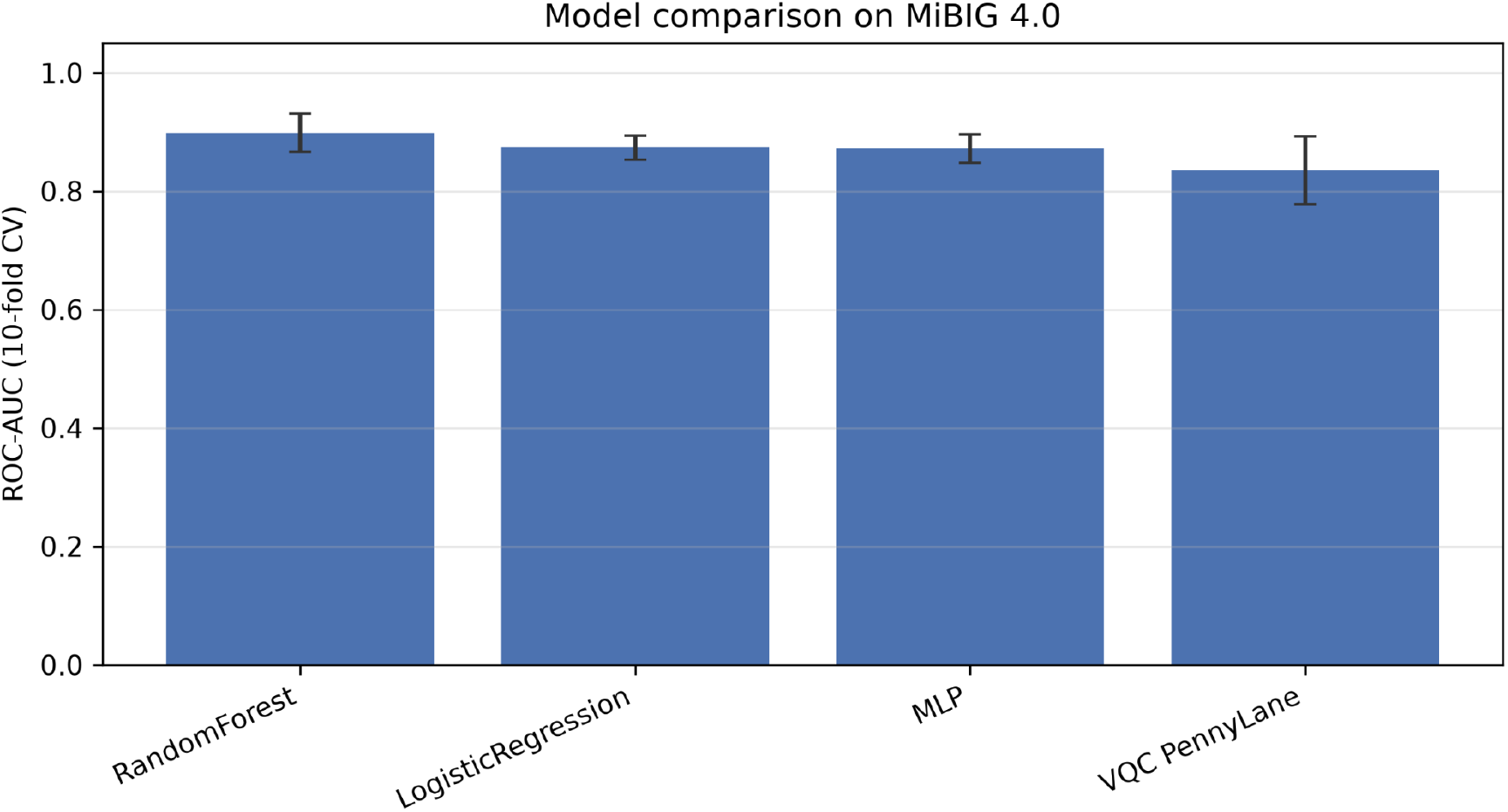
Bar chart of ROC-AUC across models.

**Figure 3:**
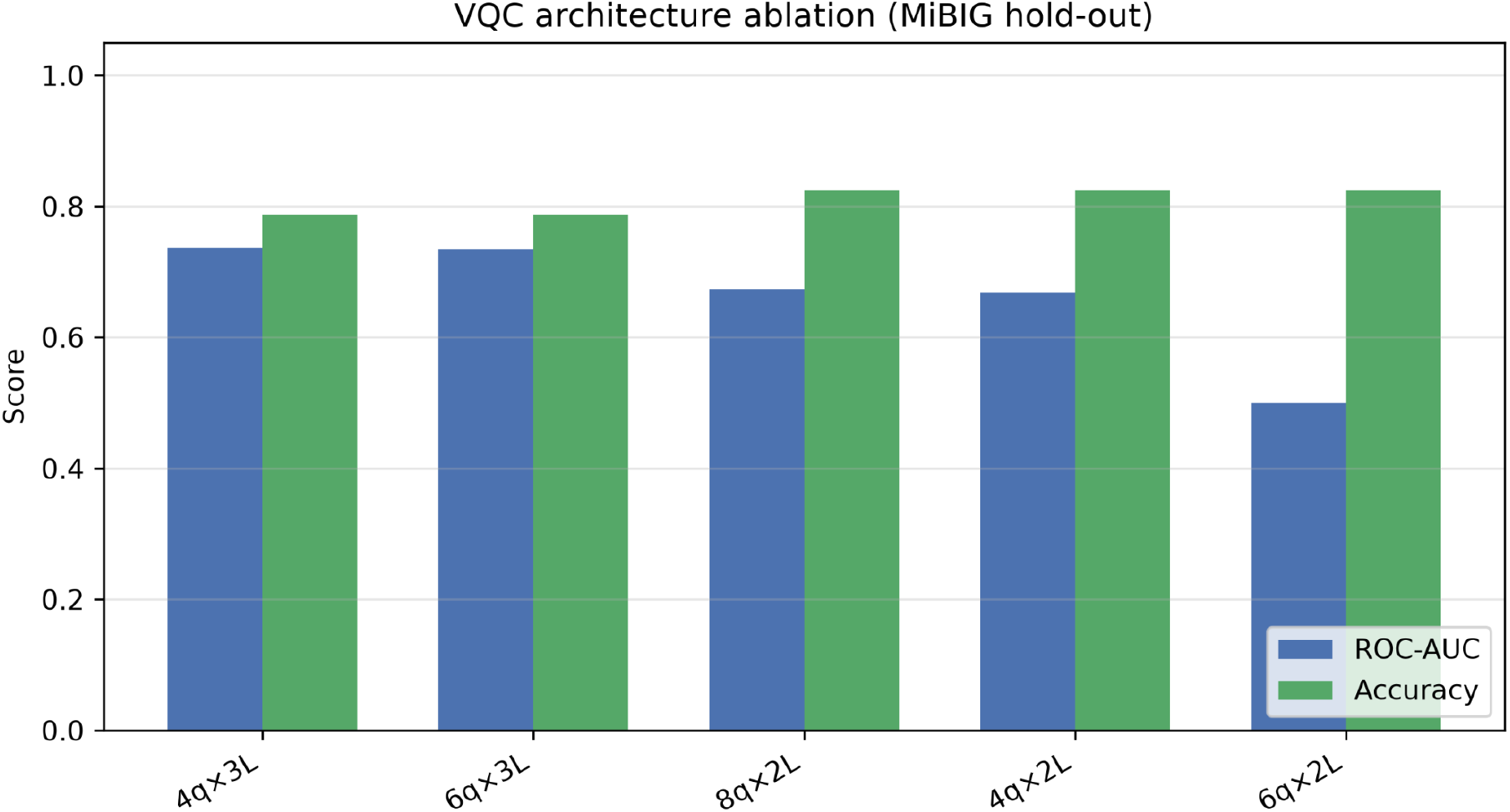
Architecture ablation chart.

**Figure 4:**
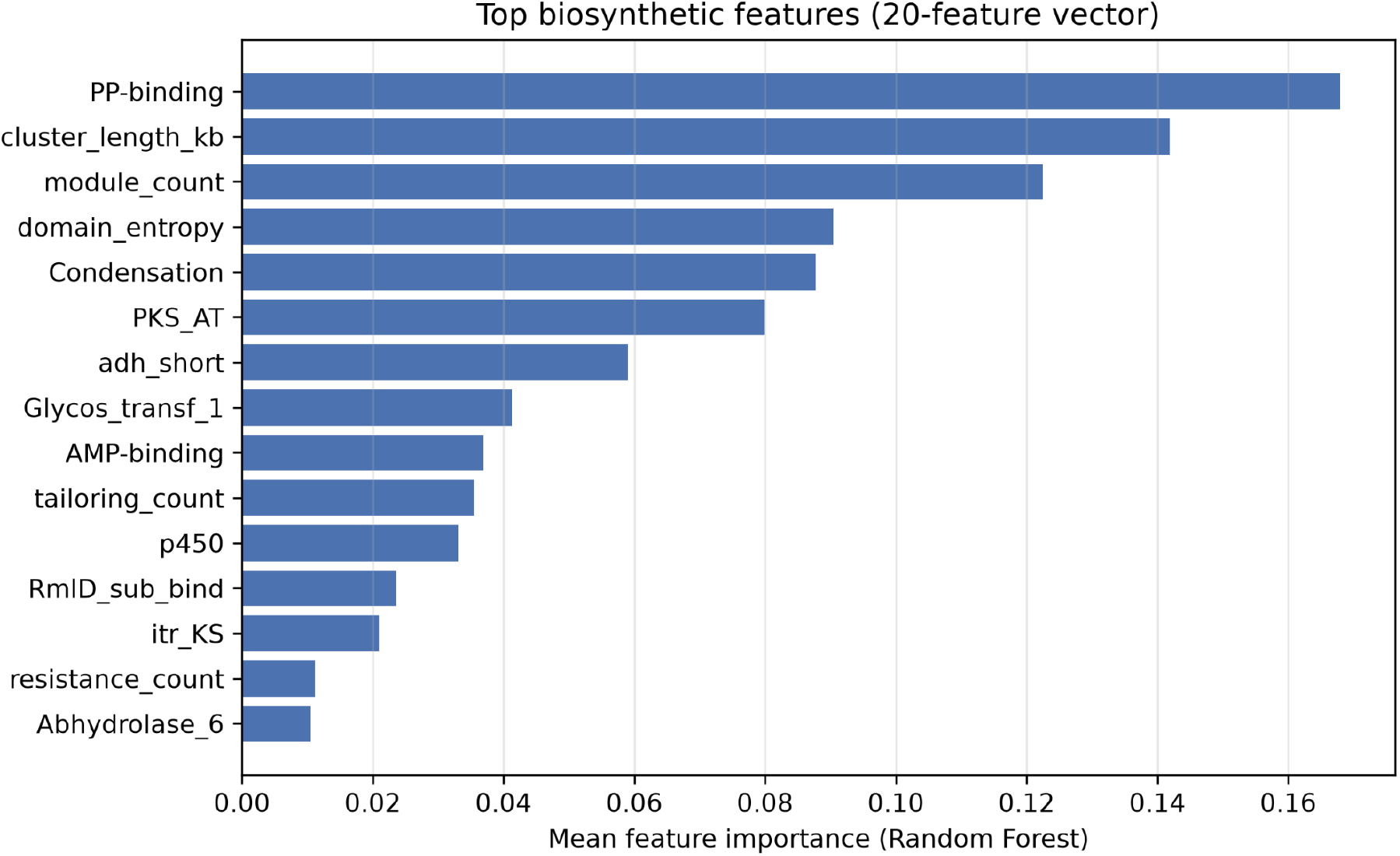
Feature importance bar chart.

**Figure 5:**
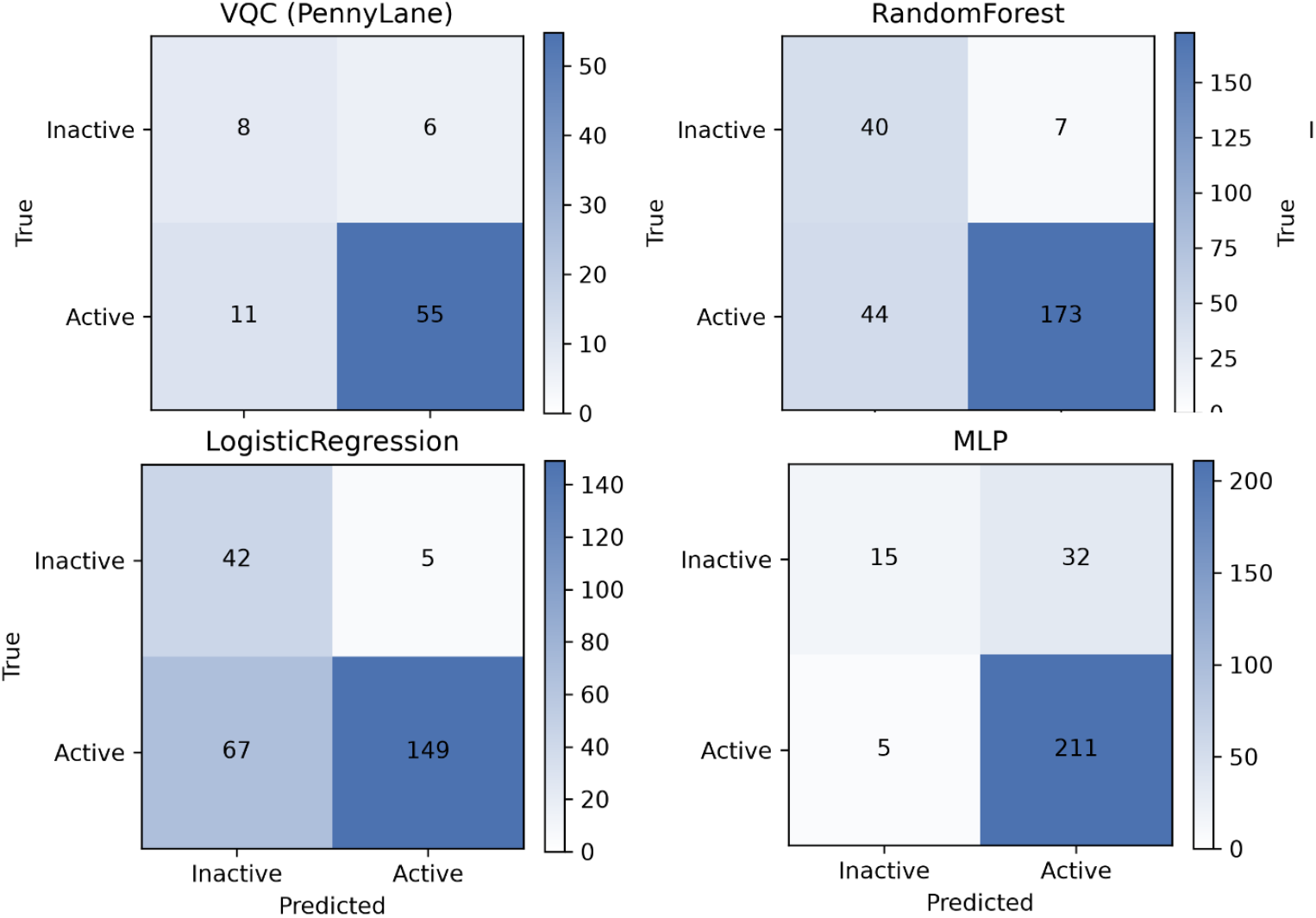
Confusion matrices.

**Figure 6:**
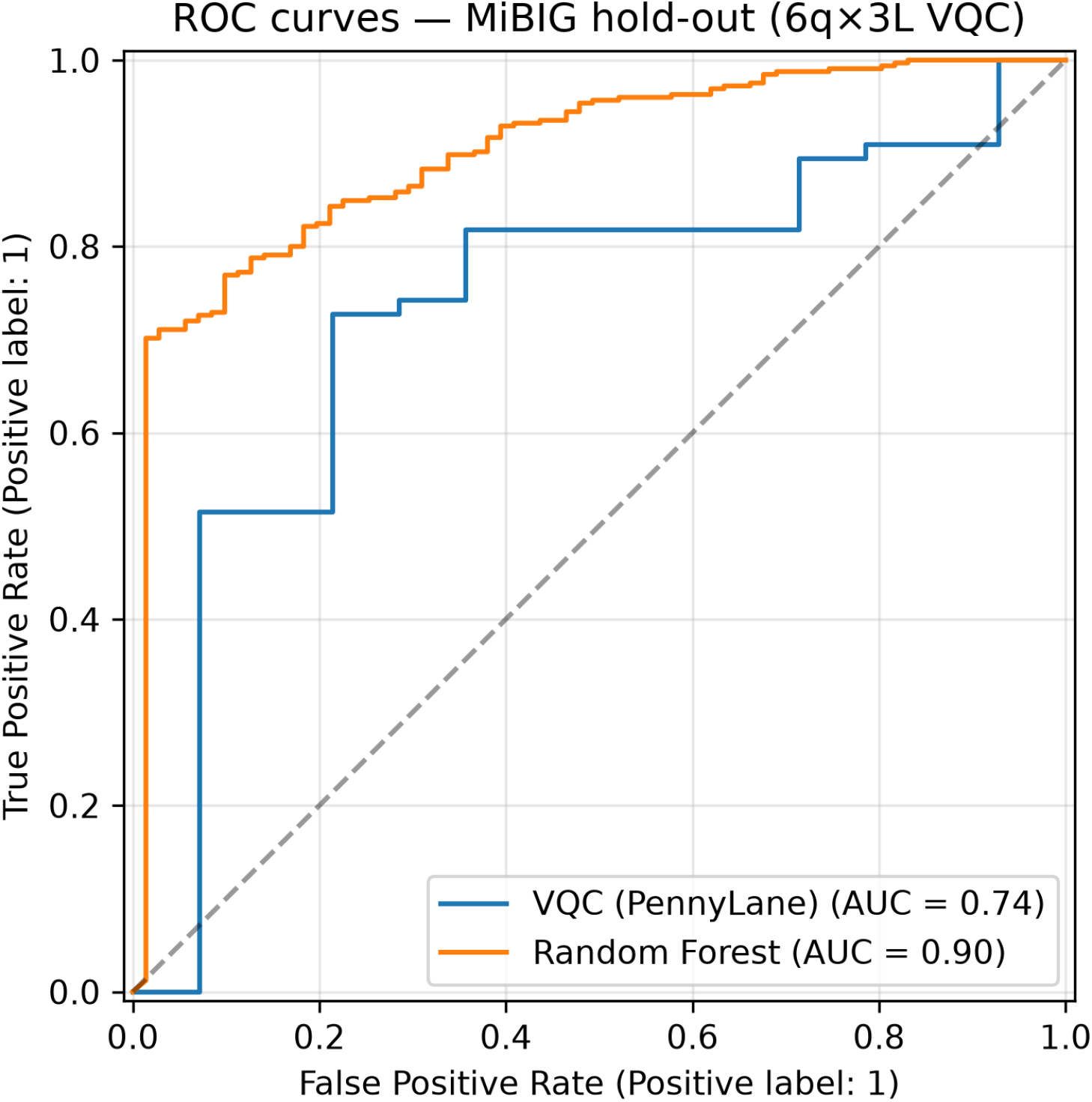
Hold-out ROC curves for VQC and Random Forest.

**Figure 7:**
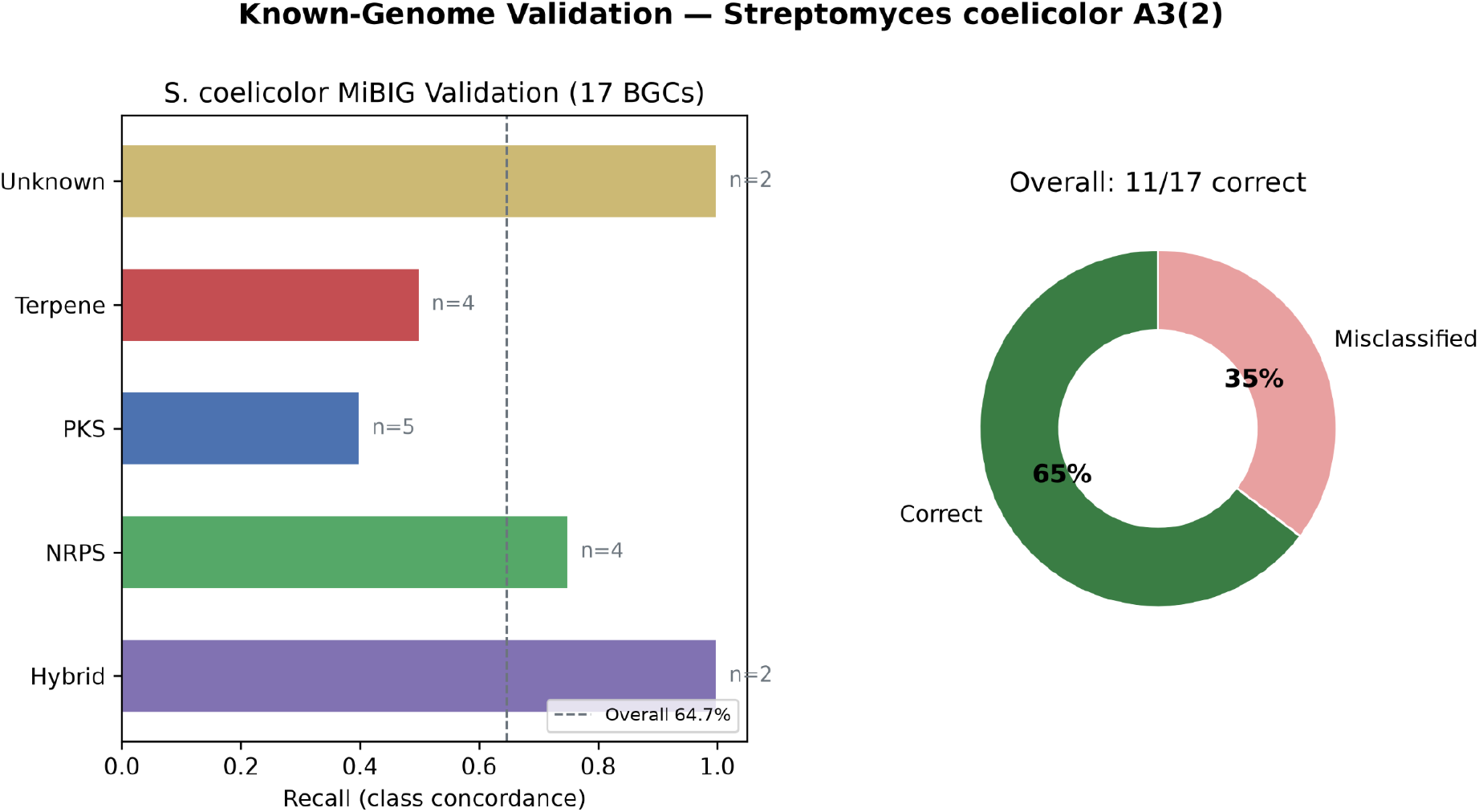
Known-genome validation on 17 *S. coelicolor* MiBIG BGCs: per-class recall and overall concordance.

**Figure 8:**
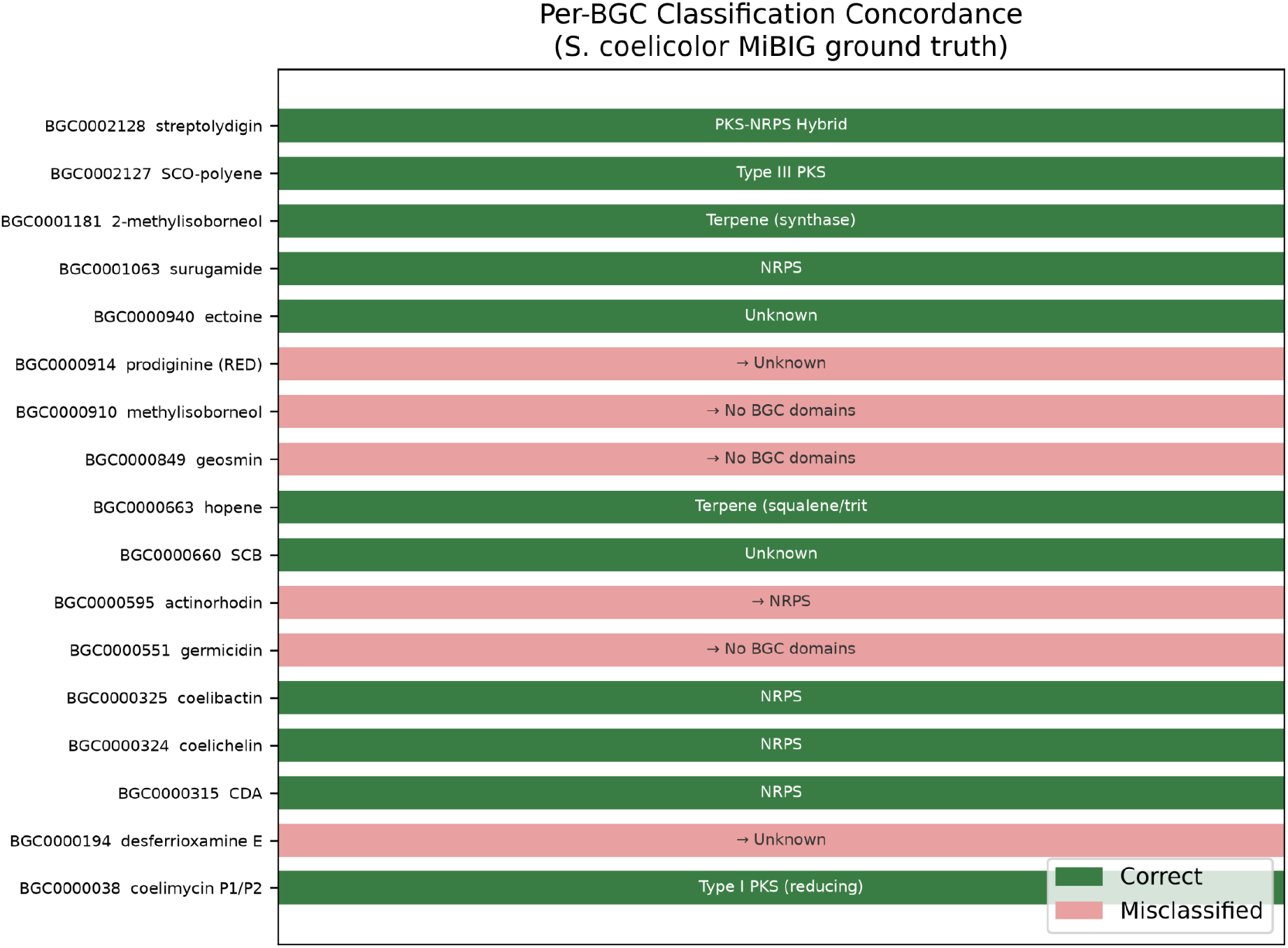
Per-BGC classification concordance matrix.

**Figure 9:**
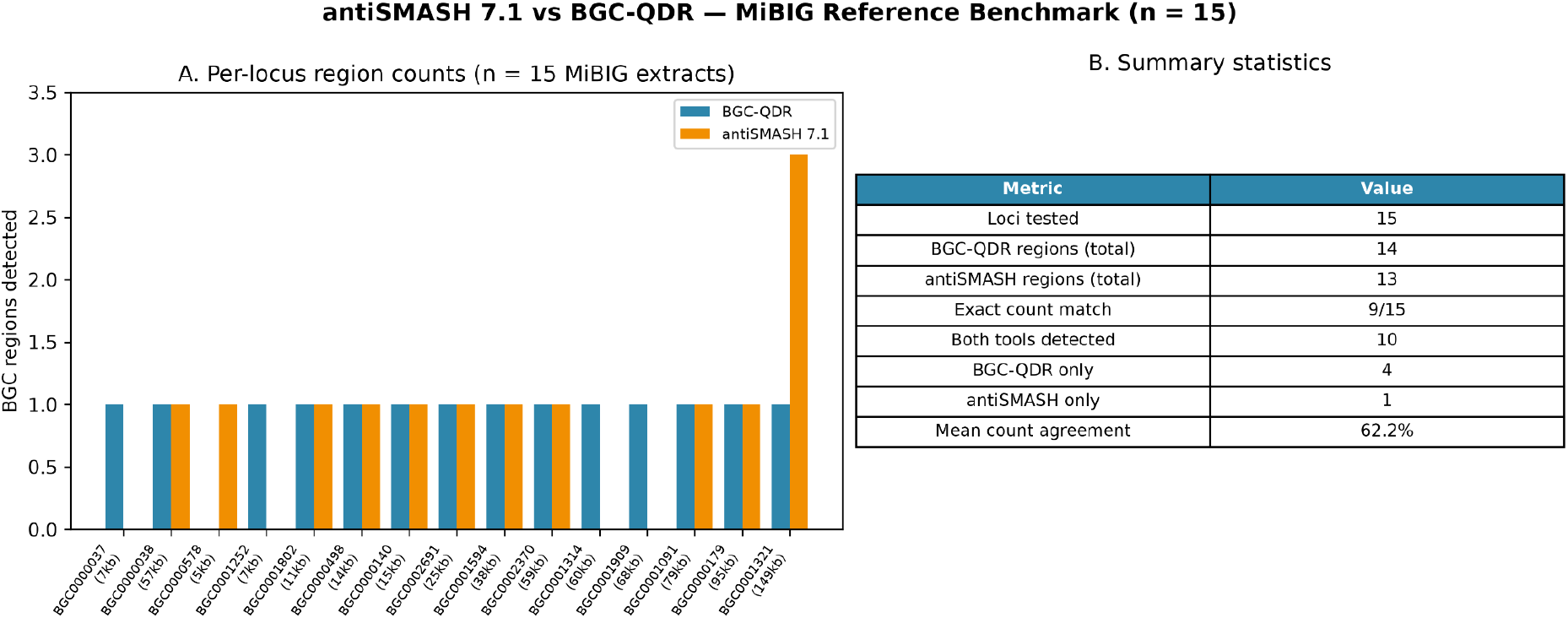
antiSMASH 7.1 vs BGC-QDR region detection on 15 MiBIG nucleotide extracts.

**Figure 10:**
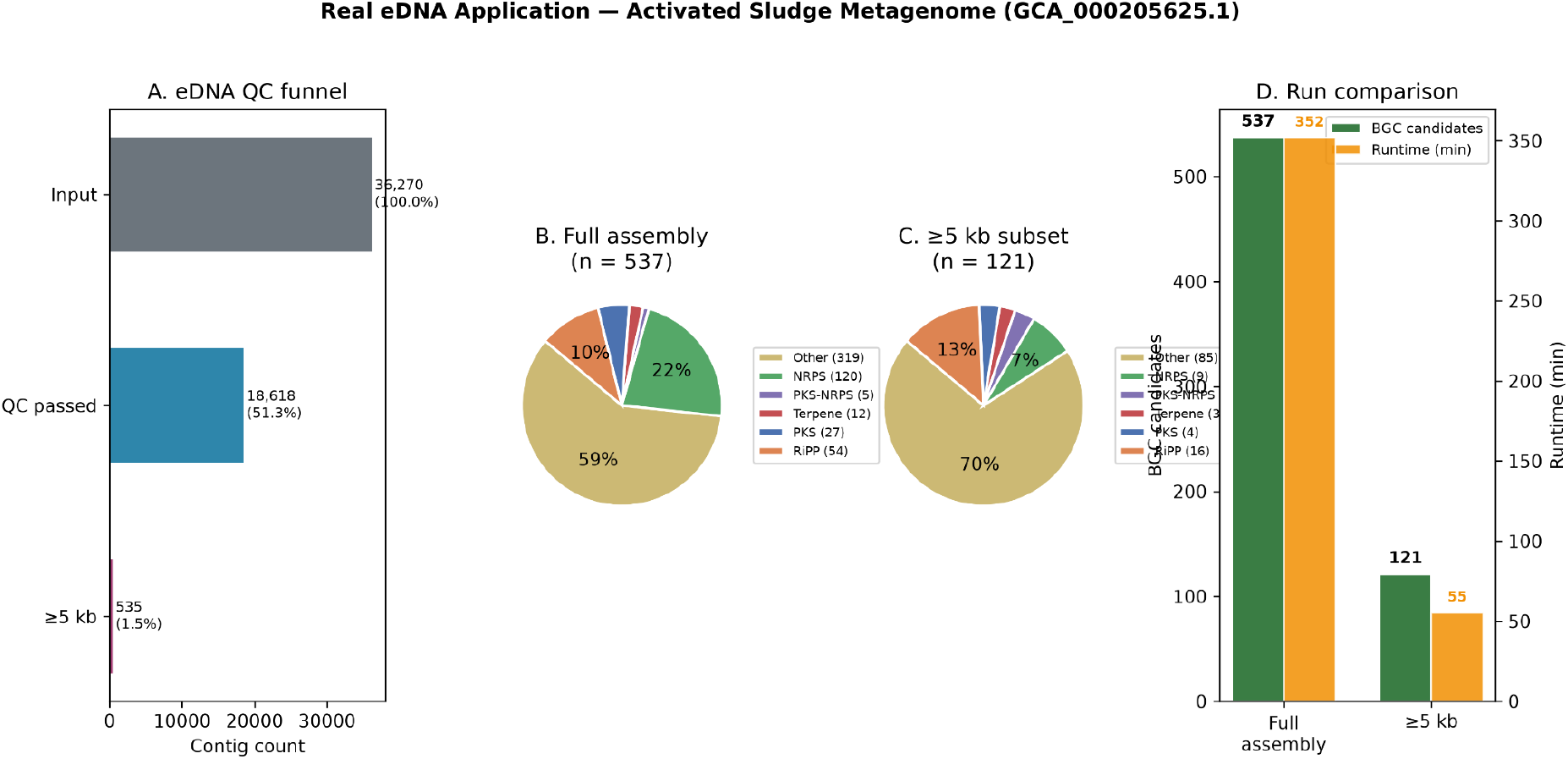
Real eDNA application on activated-sludge metagenome: QC funnel, full-assembly vs ≥5 kb BGC class breakdown, and run comparison.

**Figure 11:**
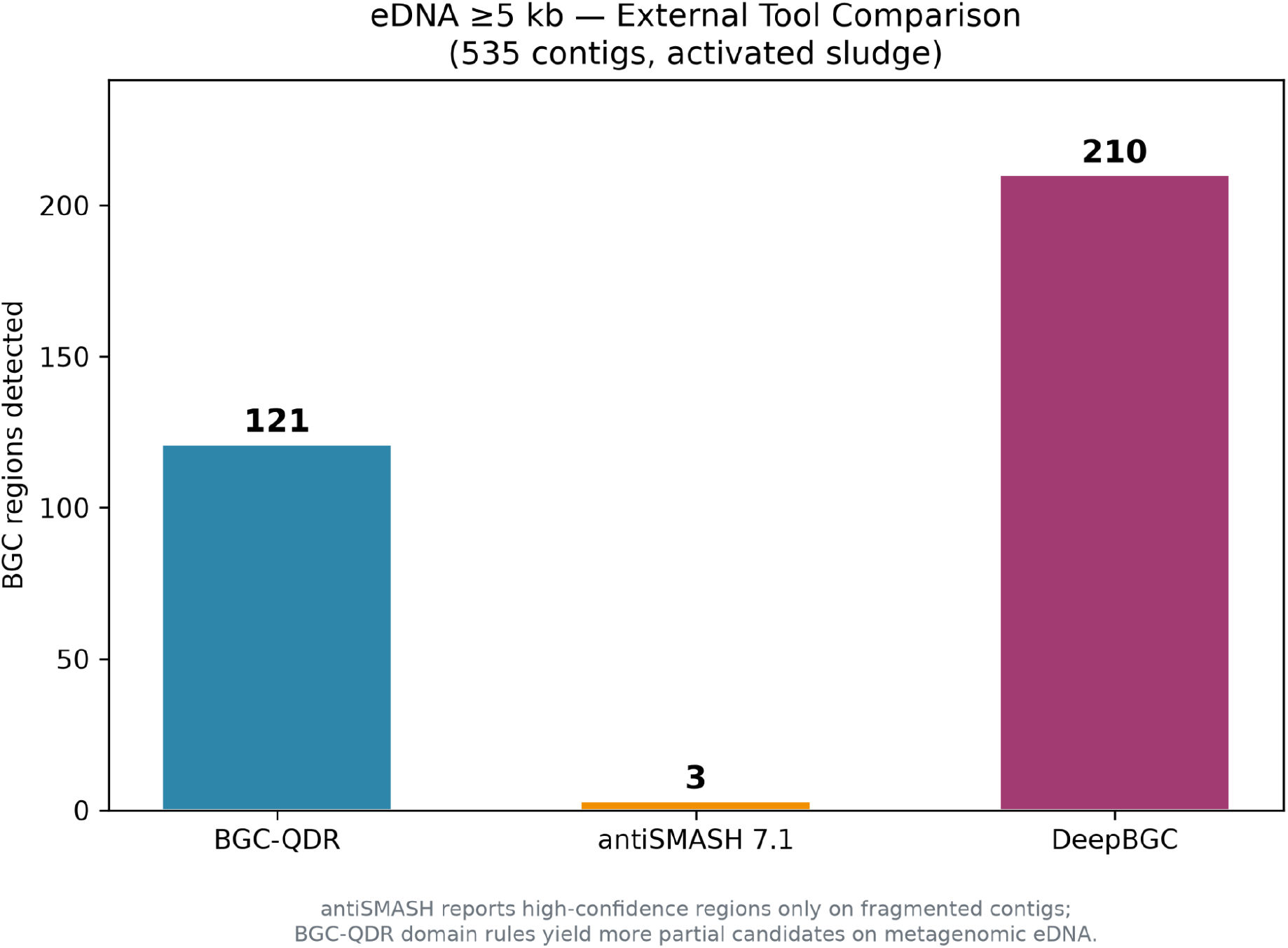
BGC-QDR vs antiSMASH vs DeepBGC on eDNA ≥5 kb.

- **Software:** Python 3.10+, PennyLane 0.45, scikit-learn, Prodigal, HMMER; experiments on CPU (PyTorch 2.12).

## IV. Results

### A. Cross-Validation Performance (Table I)

Stratified 10-fold cross-validation on MiBIG 4.0 yields the following results:

#### Key findings

- Random Forest achieves the **highest ROC-AUC** (0.898), outperforming VQC by 0.063 AUC points on average.
- MLP achieves the **highest accuracy** (0.857) and **recall** (0.976), with competitive ROC-AUC (0.872).
- VQC achieves moderate **precision** (0.910) and competitive **F1** (0.860), but trails ensembles on ROC-AUC.

### B. Statistical Significance (Table II)

**TABLE II.** VQC vs classical baselines (Wilcoxon signed-rank, 10 folds)

| Comparison | *p*-value | Significant ($\alpha = 0.05$) |
| --- | --- | --- |
| VQC vs Random Forest | 0.0098 | **Yes** (VQC lower AUC) |
| VQC vs Logistic Regression | 0.037 | **Yes** (VQC lower AUC) |
| VQC vs MLP | 0.064 | No |

Wilcoxon signed-rank tests on per-fold ROC-AUC scores:

With ten paired folds, Wilcoxon tests have adequate resolution to detect fold-wise AUC differences. VQC ROC-AUC is **significantly lower** than Random Forest and Logistic Regression, consistent with the effect sizes in Table I. The comparison versus MLP is borderline (*p* = 0.064). These results support positioning classical ensembles as stronger rankers on this benchmark, while VQC remains competitive on accuracy-oriented metrics at the default threshold.

### C. Architecture Ablation (Table III)

**TABLE III.** VQC architecture ablation (sorted by ROC-AUC)

| Config | Parameters | PCA variance | Accuracy | ROC-AUC | Time (s) |
| --- | --- | --- | --- | --- | --- |
| **4q × 3L** | 36 | 66.8% | 0.787 | **0.737** | 186.9 |
| 6q × 3L | 54 | 78.4% | 0.787 | 0.735 | 257.0 |
| 8q × 2L | 48 | 87.4% | 0.825 | 0.673 | 231.5 |
| 4q × 2L | 24 | 66.8% | 0.825 | 0.669 | 144.9 |
| 6q × 2L | 36 | 78.4% | 0.825 | 0.500 | 190.5 |

**TABLE IV.** *S. coelicolor* MiBIG class concordance (17 BGCs)

| Metric | Result |
| --- | --- |
| Correctly classified | 11 / 17 (64.7%) |
| antiSMASH literature concordance | 11 / 17 (64.7%) |
| NRPS sensitivity | 3 / 4 (75.0%) |
| PKS sensitivity | 2 / 5 (40.0%) |
| Terpene sensitivity | 2 / 4 (50.0%) |

Hold-out validation on MiBIG (250 train / 80 validation subset):

The best hold-out configuration is **4 qubits × 3 layers** (36 parameters). Increasing qubits beyond 4 does not improve ROC-AUC; deeper circuits (3 layers) outperform 2 layers at matched qubit count.

### D. Feature Importance

Random Forest feature importance (averaged over CV folds) identifies:

1. **PP-binding** (0.168 ± 0.006): peptidyl carrier protein domains
2. **cluster_length_kb** (0.142 ± 0.005)
3. **module_count** (0.122 ± 0.003)
4. **domain_entropy** (0.091 ± 0.002)
5. **Condensation** (0.088 ± 0.005)

Carrier protein and cluster architecture features dominate, consistent with modular PKS/NRPS biology.

### E. Confusion Matrices and ROC Curves

Average 10-fold confusion matrices show MLP maintains very high recall for active BGCs, while Random Forest achieves better inactive-class specificity.

### F. Known-Genome Validation (*S. coelicolor*)

We validated domain scanning and BGC class prediction against 17 curated MiBIG 4.0 entries from *Streptomyces coelicolor* A3(2). Proteins were extracted from each GBK and scanned with 51 BGC-relevant Pfam HMMs (pyhmmer).

Misclassifications concentrated on Type II PKS (actinorhodin), Type III PKS (germicidin), small terpene clusters (geosmin, methylisoborneol), and NRPS-independent siderophores (desferrioxamine E). These reflect known HMM coverage gaps rather than pipeline failure on well-represented modular PKS/NRPS loci (e.g., coelimycin, CDA, surugamide: all correct).

### G. MiBIG antiSMASH Benchmark (n = 15)

We compared BGC-QDR and antiSMASH 7.1.0 on **15 curated MiBIG 4.0 nucleotide extracts** (5–150 kb): BGC0000037, BGC0000038, BGC0000578, BGC0001252, BGC0001802, BGC0000498, BGC0000140, BGC0002691, BGC0001594, BGC0002370, BGC0001314, BGC0001909, BGC0001091, BGC0000179, and BGC0001321. The set includes the original coelimycin and erythromycin fragment case studies plus a length-stratified sample. antiSMASH was run via Docker (antismash/standalone:7.1.0, Prodigal gene finding). This supplements, but does not replace, the metagenomic tool comparison in Section IV.I (Table VII).

Representative per-locus outcomes include **BGC0000038** (coelimycin, 57,887 bp): both tools call 1 region (BGC-QDR: PKS-NRPS; antiSMASH: T1PKS). **BGC0000037** (erythromycin fragment, 7,821 bp): BGC-QDR calls 1 partial PKS region; antiSMASH calls 0 (fragment too short). **BGC0001321** (149,366 bp): BGC-QDR calls 1 Terpene region; antiSMASH calls 3 terpene subregions. **BGC0000578** (5,005 bp): antiSMASH calls 1 lassopeptide region; BGC-QDR calls 0 (domain score below threshold).

Overall, BGC-QDR is slightly more sensitive on short or marginal loci (4 BGC-QDR-only calls vs 1 antiSMASH-only), while antiSMASH can subdivide large loci into multiple regions. Full per-locus results are in benchmark_results/mibig_antismash_benchmark/mibig_antismash_report.json.

### H. Real eDNA Application

Applied to activated-sludge metagenome GCA_000205625.1 (36,270 contigs; NCBI assembly AERA01):

**BGC class distribution (full assembly v2, n = 537):** Other 319, NRPS 120, RiPP 54, PKS 27, Terpene 12, PKS-NRPS 5. All candidates are partial completeness on fragmented contigs.

**BGC class distribution (≥5 kb, n = 121):** Other 85, RiPP 16, NRPS 9, PKS 4, PKS-NRPS 4, Terpene 3.

**VQC ranking on applied eDNA:** VQC was run separately on the ≥5 kb v2 candidates (pipeline detection used --skip-vqc). All 121 candidates were scored with the MiBIG-trained 6q×3L VQC (MiBIG hold-out: accuracy 0.775, ROC-AUC 0.736). Top-ranked contig: AERA01020052 (Other, final score 0.574). Contig AERA01006104 ranked NRPS (final score 0.559), which overlaps the antiSMASH NRPS-like call (Section IV.I).

## I. External Tool Comparison on eDNA (antiSMASH & DeepBGC)

The **primary external validation** of detection sensitivity compares BGC-QDR, antiSMASH 7.1.0, and DeepBGC 0.1.31 on the same ≥5 kb eDNA subset (535 contigs from GCA_000205625.1). The MiBIG benchmark (Table V, n = 15) provides additional reference-locus comparison; metagenomic behaviour is assessed here at scale.

**TABLE V.** MiBIG antiSMASH benchmark summary (n = 15 nucleotide extracts)

| Metric | BGC-QDR | antiSMASH 7.1 |
| --- | --- | --- |
| Total regions called | 14 | 13 |
| Loci with $\geq 1$ region | 14 / 15 | 13 / 15 |
| Exact count match (per locus) | 9 / 15 (60%) | — |
| Both tools detected a region | 10 / 15 | — |
| BGC-QDR only | 4 loci | — |
| antiSMASH only | 1 locus | — |
| Mean per-locus count agreement | 62.2% | — |

**TABLE VI.** eDNA pipeline runs.

| Run | Input | QC passed | ORFs | BGC candidates | VQC ranked | Runtime |
| --- | --- | --- | --- | --- | --- | --- |
| Full assembly (v1) | 36,270 contigs | 18,618 (51.3%) | 35,759 | 0* | NA | 38.5 min |
| ≥5 kb subset (v2, batched hmmscan) | 535 contigs | 535 (100%) | 5,617 | **121** | **121** | 55.3 min |
| Full assembly (v2, batched hmmscan) | 36,270 contigs | 18,618 (51.3%) | 35,759 | **537** | NA | **351.9 min** |

\*v1 failed at domain annotation because single-pass Pfam scanning timed out on 35,759 proteins; v2 uses batched scanning (250 proteins per batch).

DeepBGC was installed using the official Conda workflow (Python 3.7, HMMER, Prodigal, DeepBGC 0.1.31 with --prodigal-meta-mode). Runtime was 23.4 min on 535 contigs.

antiSMASH called 3 regions on 535 metagenomic contigs compared with 121 BGC-QDR candidates and 210 DeepBGC regions. antiSMASH is the most conservative; DeepBGC’s neural detector is the most sensitive on fragmented contigs; BGC-QDR sits between them with interpretable domain rules plus VQC ranking. Contig AERA01006104 shows direct overlap: BGC-QDR NRPS (partial) and antiSMASH NRPS-like region agree.

## V. Discussion

### A. Quantum-Assisted Ranking

Our results demonstrate that a VQC can be integrated into a practical BGC discovery pipeline and achieves moderate **F1** (0.860) and **precision** (0.910) on MiBIG 4.0. However, **Random Forest consistently achieves higher ROC-AUC** (0.898 vs 0.835), a gap confirmed by Wilcoxon testing (Table II). We therefore position BGC-QDR as a **quantum-assisted ranking framework** that combines biologically grounded detection with a hybrid quantum-classical scorer, not as evidence of quantum computational advantage.

Taken together, these results highlight three practical strengths of BGC-QDR:

1. **End-to-end integration:** detection, novelty, and ranking in one reproducible workflow.
2. **Biological interpretability:** domain-driven features with clear importance rankings.
3. **QML feasibility:** VQC training on biosynthetic features converges and achieves moderate F1 (0.860) on MiBIG, even when ROC-AUC trails ensembles.

### B. Why Classical Models Win on AUC

Random Forest excels on tabular biosynthetic features with moderate sample size (2,636 BGCs) and class imbalance. Gradient-boosted and ensemble methods are well-suited to this regime. The VQC’s PCA compression (20 → 6 features for 6 qubits) discards variance that may be important for probability calibration, which likely contributes to lower ROC-AUC (0.835) despite moderate recall (0.831) at the default 0.5 threshold.

### C. Architecture Ablation Insights

Smaller circuits (4q × 3L) generalize better on hold-out data than larger configurations (8q × 2L). This aligns with barren plateau and overfitting concerns in deeper VQC ansätze on small biological datasets. Future work should explore 4q × 3L as the production default.

### D. Feature Biology

Dominance of PP-binding, cluster length, and module count confirms that modular PKS/NRPS architecture, not single-domain presence, drives drug-potential classification. This supports the biological validity of the feature engineering approach independent of the choice of classifier.

### E. Class Imbalance and Metrics

The 82/18 active/inactive split (Section III.C) partly explains why models differ in accuracy, precision, and recall. MLP achieves the highest accuracy (0.857) with very high recall (0.976) but moderate precision, consistent with majority-class and recall-heavy behaviour. Random Forest achieves the best ROC-AUC (0.898) and the highest precision (0.967) but lower recall (0.731), reflecting stronger specificity for inactive BGCs. VQC trails on ROC-AUC (0.835) with moderate recall (0.831) and high variance across folds. **ROC-AUC is therefore the fairest single metric for ranking models** under imbalance; raw accuracy alone favours majority-class prediction.

### F. Statistical Significance of Fold-Level Tests

Ten-fold Wilcoxon comparisons (Table II) resolve the power limitation of five-fold evaluation: VQC ROC-AUC is significantly lower than Random Forest (*p* = 0.0098) and Logistic Regression (*p* = 0.037). The MLP comparison is not significant at α = 0.05 (*p* = 0.064). Together with Table I effect sizes, these tests support treating classical ensembles as stronger rankers on MiBIG 4.0 rather than statistically equivalent alternatives to the VQC.

### G. External Tool Comparison on Metagenomic eDNA

antiSMASH and DeepBGC comparisons on eDNA (Table VII) remain the primary evidence for metagenomic detection behaviour: antiSMASH is conservative on short contigs (3 regions on 535 contigs); DeepBGC is most sensitive (210 regions); BGC-QDR is intermediate (121 regions). Contig AERA01006104 shows overlap between BGC-QDR (NRPS) and antiSMASH (NRPS-like). The expanded MiBIG benchmark (Table V, n = 15) shows moderate per-locus count agreement (9/15 exact matches) with complementary failure modes: BGC-QDR-only calls on short or marginal loci, antiSMASH-only on BGC0000578, and antiSMASH region splitting on BGC0001321.

**TABLE VII.**
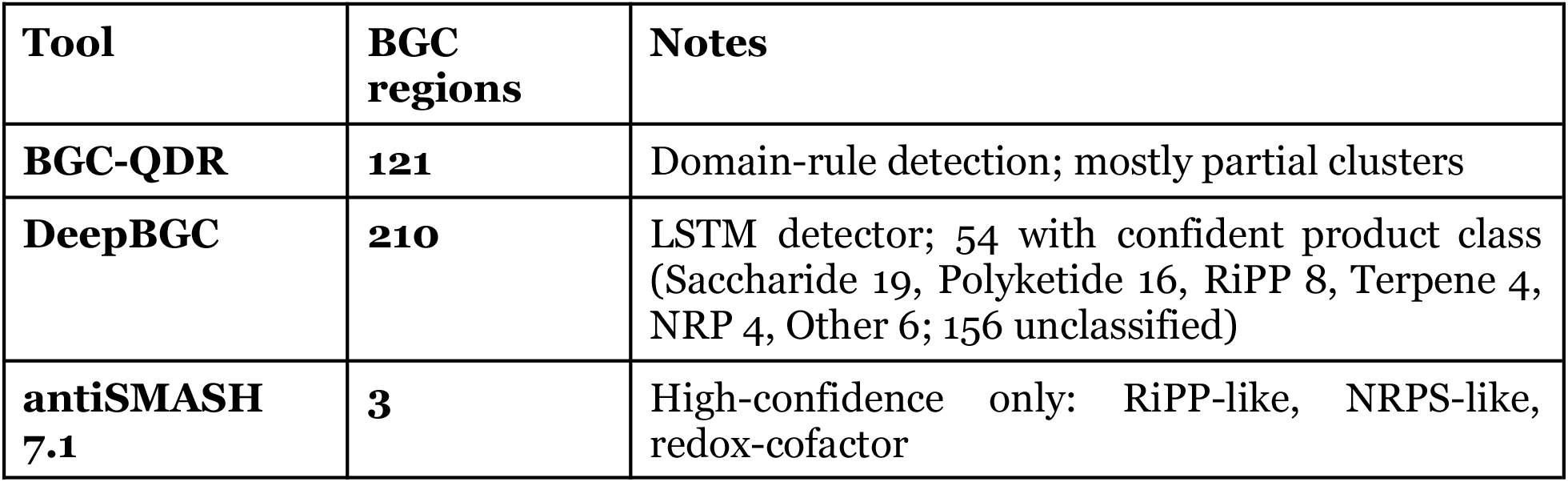
Tool comparison on eDNA ≥5 kb (535 contigs)

| Tool | BGC regions | Notes |
| --- | --- | --- |
| **BGC-QDR** | **121** | Domain-rule detection; mostly partial clusters |
| **DeepBGC** | **210** | LSTM detector; 54 with confident product class (Saccharide 19, Polyketide 16, RiPP 8, Terpene 4, NRP 4, Other 6; 156 unclassified) |
| **antiSMASH 7.1** | **3** | High-confidence only: RiPP-like, NRPS-like, redox-cofactor |

## H. Limitations

1. **No quantum advantage demonstrated:** classical ensembles outperform VQC on ROC-AUC, with statistically significant gaps versus Random Forest and Logistic Regression (Table II).
2. **Simulator only:** all quantum experiments use PennyLane statevector simulation; hardware noise not evaluated.
3. **MiBIG bias:** training data over-represents actinobacterial, clinically studied BGCs; labels are coarse active/inactive proxies.
4. **MiBIG antiSMASH benchmark scope:** 15 reference loci (not exhaustive MiBIG coverage); region-count agreement is moderate (60% exact match).
5. **eDNA fragmentation:** short contigs may yield incomplete domain architectures; detection sensitivity depends on contig length. Large metagenomes require batched domain scanning (default batch size: 250 proteins). VQC ranking was applied to the ≥5 kb eDNA subset only (121 candidates), not the full 537-candidate assembly run.
6. **Binary labels:** MiBIG active/inactive is a coarse proxy for true drug potential.

## VI. Conclusion

We presented BGC-QDR, an open-source pipeline for biosynthetic gene cluster discovery and quantum-assisted drug-potential ranking from environmental DNA. The workflow integrates quality control, Prodigal ORF prediction, Pfam HMM annotation, rule-based BGC classification, MiBIG novelty assessment, and PennyLane VQC ranking into a single reproducible tool.

On MiBIG 4.0 (2,636 BGCs), stratified 10-fold cross-validation shows VQC accuracy of 0.789 ± 0.076 and ROC-AUC of 0.835 ± 0.057, with Random Forest achieving the strongest ROC-AUC (0.898 ± 0.032). Wilcoxon tests confirm significantly lower VQC AUC versus Random Forest (*p* = 0.0098) and Logistic Regression (*p* = 0.037). A 15-locus MiBIG antiSMASH benchmark (Table V) and metagenomic tool comparison (Table VII) provide external validation. Architecture ablation favors 4 qubits × 3 layers for VQC hold-out performance.

BGC-QDR demonstrates that quantum-assisted ranking can be embedded in a real bioinformatics pipeline, but **classical ensemble methods remain the stronger ranker on this benchmark**. Future work will evaluate hardware-deployed VQCs, expanded eDNA cohorts, and regression on quantitative bioactivity data. Source code, reproduction instructions, and benchmark figures are available in the project repository.

